# TRAP seq in 3D Angiogenesis Assays Reveals a Distinct Endothelial Translatome Associated with Early and Late Stages of Morphogenesis

**DOI:** 10.1101/2025.06.17.660138

**Authors:** Samantha King, Qing-fen Li, Ian Tschang, Ekta Singh, Ramon Bossardi Ramos, Kevin Pumiglia

## Abstract

Endothelial cells (ECs) are often a minority cell type in a tissue, limiting the utility of bulk sequencing approaches. Single cell sequencing lacks sensitivity and requires disruptive tissue digestion techniques. TRAP seq (Translating Ribosome Affinity Purification) or “RiboTag” has been used to overcome these limitations. Multicellular co-culture systems allow primary ECs to differentiate and undergo tubular morphogenesis in cell culture, however similar limitations exist with these *in vitro* assays, as ECs are under-represented by as much as a factor of 10 in many assays. We sought to use TRAP seq to better understand the gene expression landscapes that drive these morphogenic events. We found TRAP seq selectively enriches for endothelial RNA in two distinct co-culture paradigms, in both the planar and fibrin bead co-culture assays.

Intriguingly, the use of this technology in vitro, revealed distinct gene expression changes in blood vessel development and in the mitotic cell cycle, with genes unique to early and late phases of morphogenesis. It is widely accepted that expression of the NOTCH signaling pathway is a regulator of angiogenesis in vivo. Correspondingly, we found that a large number of NOTCH related genes relevant to endothelial cell biology, were changing across this morphogenic process, underpinning the importance and utility of this technology in 3D multicellular cultures that model in vivo environments.

## INTRODUCTION

Endothelial cells (ECs) comprise the innermost layer of blood vessels and are essential for the proper development and maintenance of the vasculature [1, 2]. Concentration gradients of vascular endothelial growth factor (VEGF) and the resultant signaling are responsible for the endothelial switch between the migratory tip and proliferative stalk phenotypes of angiogenic ECs [1, 2]. This signaling in the endothelium is essential for the sprouting and elongation of new blood vessels towards the site of pro-angiogenic stimuli, such as hypoxic tissues or sites of injury, where vascularization is needed, in a process called angiogenesis [1, 2]. In addition to angiogenesis, ECs are also critical for regulating the exchange of nutrients and oxygen between the bloodstream and surrounding tissues, regulation of vascular tone, hemostasis, and immune surveillance [1, 2].

Endothelial cells have traditionally been studied *in vitro* using 2D culture, however these cultures are limited by their inability to accurately mimic the complex microenvironments *in vivo,* and as a consequence, biological functions such as differentiation and tubular morphogenesis are difficult to achieve in two-dimensional culture [3]. *In vitro*, three-dimensional, multicellular systems have been described that introduce some of the *in vivo* complexities, such as interactions with mural cells and their secretome, binding and processing of extracellular matrix, and changes in biomechanical stiffness [3–5]. Importantly, these models allow for more accurate reproduction of the angiogenic process as well as disruptions in angiogenesis that can occur following altered cell signaling due to mutations or disrupted microenvironmental interactions [6–9]. However, to date there is limited data available on the transcriptional changes that accompany these *in vitro* morphogenesis events.

Bulk RNA sequencing allows expression analysis of gene changes across an entire population of cells [10]. While effective, it has limitations when studying cell types that are a minority in a population, as is often the case with vascular endothelial cells in tissues, as the RNA expression profile is dominated by non-endothelial cells. Single cell RNA seq (sc-RNAseq) is an alternative which makes libraries for sequencing from individual cells and then based on expression profiles, these individual cells are organized into clusters of distinct cell types [11].

However, this methodology also has limitations, as sc-RNA seq protocols typically involve mechanical disruption and digestion of tissues and cell aggregates. These manipulations remove the cells from their microenvironment, alter adhesion properties, and potentially introduce new signaling from the activation of protease sensitive receptors such as, PAR2, often highly expressed on endothelial cells [12]. Single cell sequencing is also limited by the low abundance of starting RNA, leading to a lack of resolution and sensitivity, particularly for low abundance transcripts.

Recently investigators studying sprouting angiogenesis and endothelial cell heterogeneity have used translating ribosome affinity purification (TRAP) sequencing to enrich endothelial cell transcript representation from various organ beds including the retina, brain, liver, and lung [13–16]. Originally developed in the Heintz lab, TRAP seq is a method for selectively isolating the actively translating mRNA from the ribosomes of a target cell type using a large ribosomal subunit epitope tag (RiboTag) [13, 15]. This allows for the immunoprecipitation of the actively translating mRNA from a specific target cell type that expresses the RiboTag from a mixed population of cells [13–16]. This strategy produces an unbiased approach in studying gene expression changes across mRNA that are actively being translated into protein for utilization by the cell [13–16]. As tissue samples can be flash frozen in real time, TRAP seq also allows for the isolation of mRNA from minority cell types in their native or non-disrupted state [13–16].

Understanding the transcriptional and translational landscapes that drive morphogenesis in normal ECs can lay a foundation for therapeutic targeting of diseases that manifest pathological angiogenesis, such as retinopathy, tumors, and vascular malformations. We sought to use TRAP seq to study changes in the endothelial translatome during 3D in vitro morphogenesis in co-culture assays. Our data reveal dynamic changes in the endothelial cell translatome following three-dimensional co-culture, particularly in genes involved in mitogenesis and blood vessel development. We also found a marked downregulation of cellular proliferation early in the process and a vast number of genes in the NOTCH signaling pathway that are being dynamically regulated throughout morphogenesis.

## Results

### Generation of a lentiviral vector harboring Rpl22-3X-HA and its use in 3D co-culture of HUVECs

Low passage primary endothelial cells infected with lentivirus maintain full differentiation capability, thus we sought to create a lentiviral vector to deliver an appropriate ribosomal tagged construct. As a commercially available mouse model (Jackson, Strain #:029977) is available and has been used with success to study ECs [14, 17], we chose to develop a Rpl22-3X-HA construct. We cloned a Rpl22-3X-HA construct into the pHAGE-IRES-tdTomato vector, generating the Rpl22-3X-HA_pHIT lentiviral vector which provides bicistronic expression of the marker gene TdTomato, providing an easy way to visualize cells during morphogenesis as well as method for drug-free selection of infected cells by flow cytometry, if necessary **(Figure 1a)**.

**Figure 1.**
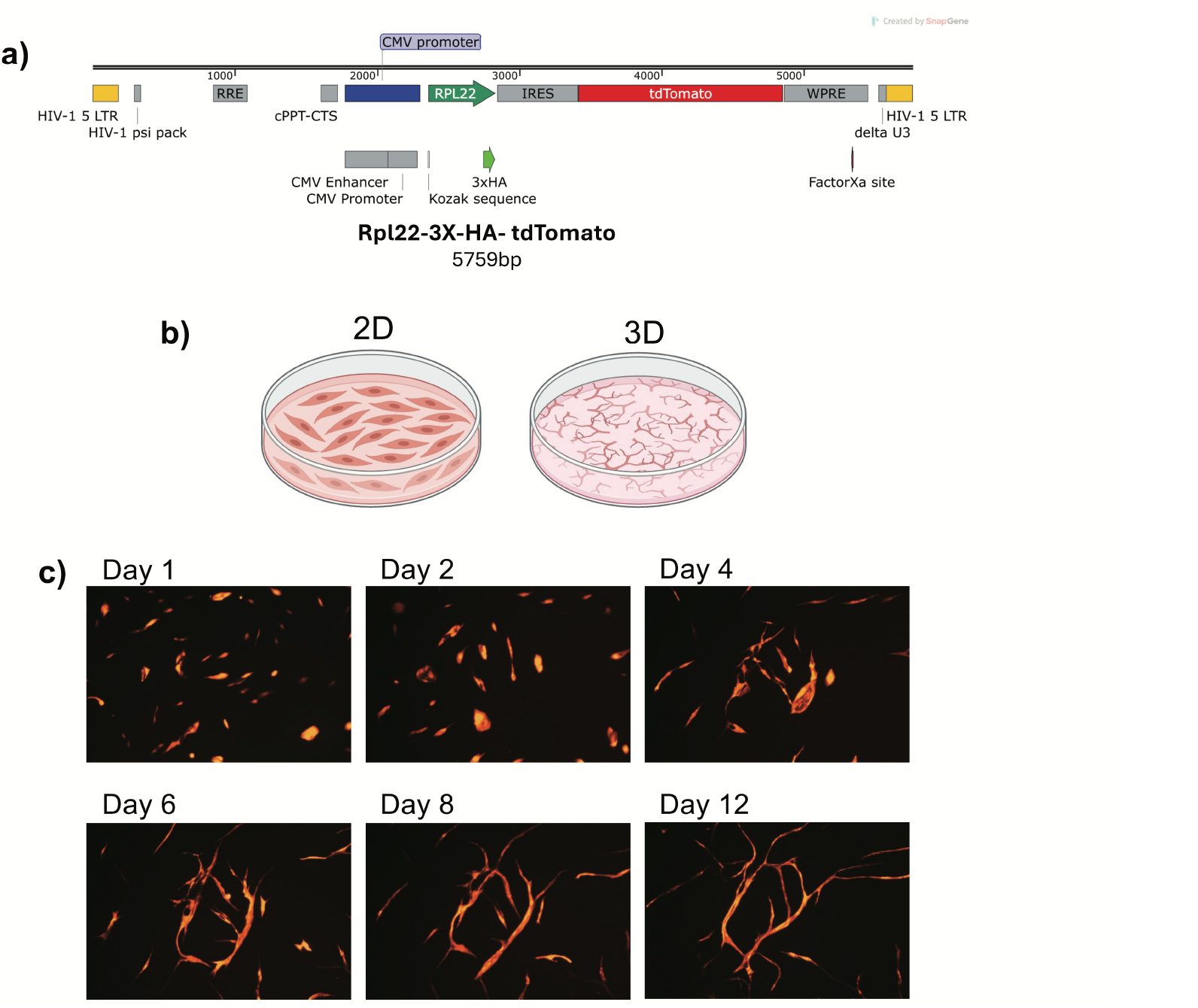
Generation of a lentiviral vector harboring Rpl22-3X-HA and its use in a 3D co-culture of HUVECs. **a)** Map of Rpl22-3X-HA-tdTomato insert used in pHAGE lentiviral backbone for use in HUVECs to be subjected to TRAP seq. Vector diagram created in SnapGene. **b)** Diagram depicting morphological differences in 2D culture of HUVECs compared to 3D culture of HUVECs cultured on top of a confluent monolayer of fibroblasts in a planar co-culture morphogenesis assay. *Created in BioRender. King, S. (2025) https://BioRender.com/s68o534*. **c)** HUVECs infected with Rpl22-3X-HA-tdTomato are co-cultured with primary fibroblasts for 12 days in the planar co-culture assay. Images are taken at 5X at the indicated time points and then equally cropped via ImageJ to highlight anastomoses. Images are representative of 4 independent experiments.

To study the changes in the endothelial translatome during cellular morphogenesis we first infected HUVECs with the Rpl22-3X-HA_pHIT vector with efficiency approaching 100%, so no further selection was used. We used an assay developed by Bishop et. al. where HUVECs are seeded atop a confluent monolayer of primary neonatal human dermal fibroblasts (NHDF) at approximately a 1:10 ratio (HUVEC:NHDF) which initiates endothelial differentiation into vessel-like structures **(Figure 1b)**. This assay is then allowed to proceed for up to 12 days by which time cells have typically formed elongated tubule-like structures that branch and anastomose (see schematic in **Figure 1b**). A time course using Rpl22-3X-HA_pHIT infected HUVECs is demonstrated in **Figure 1c**, using the td-tomato expression to visualize the endothelial cells. At day 1 of co-culture, HUVECs were well spread and appeared partially rounded and motile, migrating around the plate **(Figure 1c)**. By day 4, the HUVECs began to elongate and fuse together, which extended and continued to fuse into branching networks at day 8. By day 12 these networks showed only minimal changes and appeared quiescent **(Figure 1c).** As we and others have previously validated, many of these structures contain patent lumens [4, 8].

Due to the dramatic morphological changes that occur between days 1, 4, and 8, we decided to target those time points for RNA isolation in our TRAP seq experiments designed to better define the transcriptomic changes occurring during this morphogenic process. As biological variability is a factor when using primary cells, experiments were conducted using 4 independent pools of primary HUVECs from two different commercial vendors, each pool representing at least three donors of mixed races and sexes. To ensure representative numbers of endothelial and fibroblast mRNAs in the 2D endothelial samples, identical numbers of endothelial and fibroblasts were plated separately at the same time and media as the co-culture assays and were lysed in a common fraction of lysis buffer. A complete outline of the experimental scheme is shown in **Supplemental Figure 1** and raw counts are available in **Supplemental Table 1.**

### TRAP seq effectively enriches endothelial cell mRNA in the planar co-culture assay of morphogenesis

Co-culture samples infected with the Rpl22-3X-HA coding lentivirus were lysed and subjected to immunoprecipitation (IP) with anti-HA beads following 8 days in co-culture. These IP RNA samples were compared to input RNA samples (no immunoprecipitation), containing an enrichment with fibroblast cells by approximately 10-fold, based on plating density. As anticipated, we found significant differences in gene expression between input and IP samples **(Figure 2a).** Gene Ontology (GO) analysis of the differentially upregulated genes found strong enrichment in various types of endothelial cell signatures in the MSigDB of cell types **(Figure 2b)** as well as EC associated Biological Processes gene sets, including angiogenesis and blood vessel development **(Figure 2c).** To further validate the enrichment of ECs with TRAP, we analyzed a range of endothelial specific marker genes often employed in IHC as well as sc-RNA-seq including, VWF, SOX18, CDH5, and PECAM1. We found all to be strongly enhanced in the IP samples compared to the input **(Figure 2d)**, confirming the efficacy of this technique to study minority cell types in 3D planar co-cultures.

**Figure 2.**
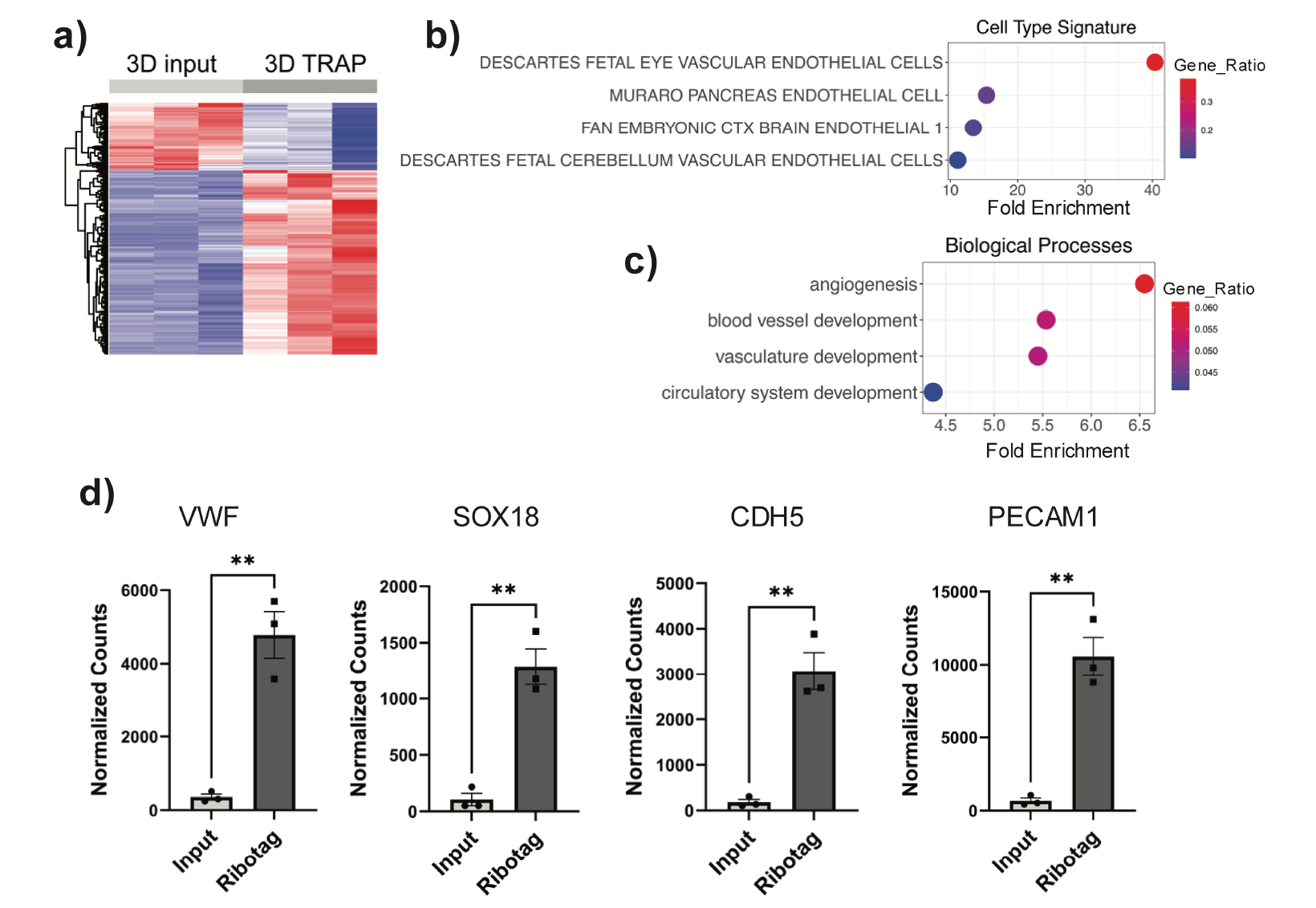
RiboTag provides significant enrichment of endothelial cell transcripts from the 3D planar co-culture. **a)** Heat map of differentially expressed genes between input and TRAP-seq samples filtered for genes changing 2-fold with p<0.05. **b)** TRAP DEGs changing increasing more than 3-fold (p <0.05), were evaluated against cell-type signatures in MSigDB, with the top terms shown. **c)** Top GO terms in biological processes following analysis of upregulated DEGs changing 3-fold (p<0.05). **d)** Normalized raw counts showing enrichment of classic EC marker genes VWF, SOX18, CDH5, and PECAM1 in TRAP samples significance determined by student’s t-test, p<0.05.

We next wanted to extend the study to another 3D *in vitro* morphogenesis assay to validate TRAP was effective in isolating RNA from a more complex and technically challenging assay. In this alternative assay, originally described by the Hughes lab [5], Cytodex beads are coated with HUVECs, followed by embedding in a fibrin gel matrix with fibroblasts seeded on top of the fibrin gel. In this environment, HUVECs differentiate and form sprouts that extend into the gel matrix **(Supplemental Figure 2a)**. We validated the endothelial enrichment with TRAP seq in this assay **(Supplementary Figure 2b-e, Supplemental Table 2)** extending the utility of this technology in other complex organoid systems.

### TRAP seq of 3D co-culture of HUVECs reveals distinct biological processes involved with morphogenesis over time

The principal objective of these studies is to investigate the change in EC gene expression in 3D co-cultures that are permissive for morphogenesis compared to traditional 2D culture of HUVECs. We found HUVECs cultured alone in 2D versus cultured in the 3D model with fibroblasts, had a substantial number of differentially expressed genes after 24 hours **(Supplemental Figure 3a and 3b)**. Confirming what we observed visually in the co-cultures, gene ontology analysis of DEGs found various biological processes changing with 3D culture such as, cell migration and vasculature development **(Supplemental Figure 3c)**. To this point, we observed a number of genes (e.g. CXCR4, KDR, DLL4, and UNC5B) critical for migration and tip cell specification enriched at this early time point **(Supplemental Figure 3d**), while other genes, showed marked down regulation, including SOX4, JAG1, CCN2, and PDGFB **(Supplemental Figure 3e)**. These data demonstrate that after just 24 hours in co-culture, HUVECs are altering their transcriptome in ways consistent with angiogenesis and blood vessel morphogenesis, underscoring the importance of modeling endothelial biology in complex 3D environments.

We next sought to determine which genes are important for regulating morphogenesis of HUVECs into vessel-like structures from the beginning stages of differentiation to maturation.

Principal component analysis from all samples included in the RNA seq dataset revealed unique clustering at each time point, and that gradual changes in gene expression occur as HUVECs progress through morphogenesis **(Figure 3a)**. This gradual change and unique clustering suggested that there might be sets of genes that are uniquely altered at different stages of morphogenesis. We determined the differentially expressed genes (DEGs) for each time point by comparing each 3D time point to the same control 2D dataset. This allowed us to determine which individual DEGs and biological processes are changing at each time point and which ones are shared, when compared to a common control. GO analysis of DEGs from early to late morphogenesis showed unique upregulated and downregulated genes at each time point (Supporting data table; Main-3B). Interestingly, across the time points two Biological Processes in particular were enriched in nearly all circumstances, *Blood Vessel Development* and *Mitotic Cell Cycle* **(Figure 3b)**. Further analysis of the GO term *Blood Vessel Development* revealed the genes have a balance of up and down regulated genes, that gradually increases in the number as you progress through morphogenesis **(Figure 3b)**. However, the genes that were changing at each time point were distinct (Supporting data table; Main 3B top GO DEGs). The *Mitotic Cell Cycle* biological process was absent from the analysis at the day 1 point but was highly enriched by day 4 and is maintained throughout day 8 of the co-culture **(Figure 3b)**.

**Figure 3.**
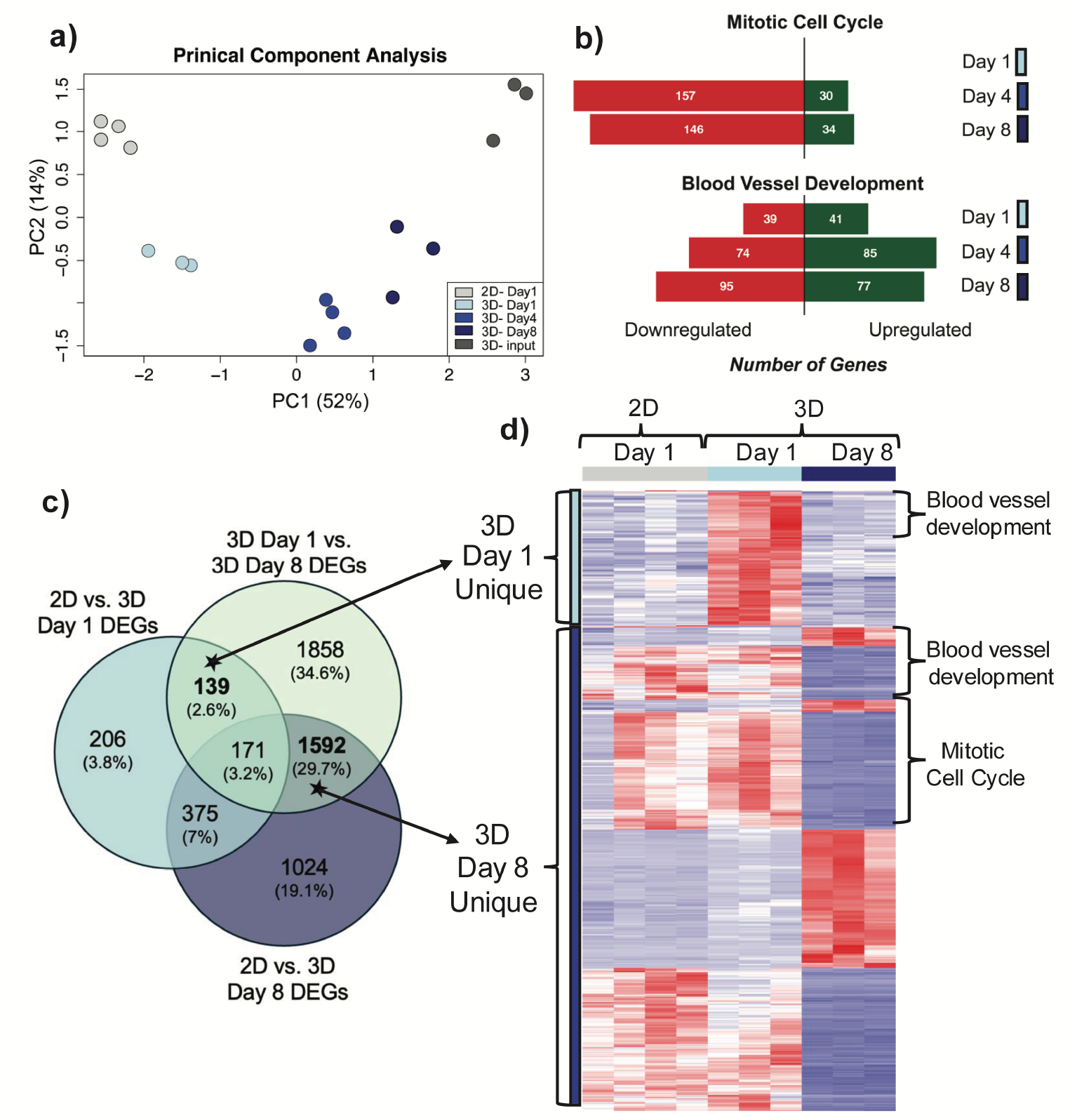
RiboTag reveals distinct biological processes involved with normal vascular morphogenesis. **a)** PCA plot of groups included in TRAP seq analysis. **b)** Top GO terms from Metascape analysis (p-value <0.05 and logFC >1 and logFC< −1) of DEGs between 2D day 1 and 3D day 1, or 3D day 4, or 3D day 8. The number of individual gene hits for that GO term that are upregulated are shown in green and downregulated are shown in red. **c)** Venn diagram from set analysis between 2D vs. 3D day 1 (light blue), 2D vs. 3D day 8 (dark blue), and 3D day 1 vs. 3D day 8 (gray). This comparison allowed for the selection of unique genes that are important for 3D culture at early or late time points. **d)** Genes unique to days 1 and 8 were combined and clustered to represent by a Heatmap. Genes derived from the blood vessel development and mitotic cell cycle GO term are highlighted. Day 1 unique genes are denoted by the light blue bars, and the day 8 unique genes by the dark blue bars (DEGs with Log FC +/− 1, p-value <0.05).

Other terms that were highly enriched included processes related to *cell migration* at day 1 and *extracellular matrix remodeling* and *cellular metabolism* at day 8. Despite the commonality in the GO terms, it was clear from both hierarchically clustered heat diagrams, as well as examination of the genes enriched in the GO terms related to blood vessel development, that there were distinct gene expression programs happening between day 1 and 8, with day 4 appearing like a transition state. To identify unique genes between the day 1 and day 8 time point, we examined genes that were different between day 1 and day 8 and then compared those genes at each time point that were different in the comparison to 2D cultures **(Figure 3c)**. Using set analysis, we found 139 genes that are unique to day 1, and 1592 genes that are unique to day 8 **(Figure 3c)**. We collated these genes, identified those related to blood vessel development or cell cycle, and clustered those in a representative heat map **(Figure 3d)** which clearly shows the unique early and late-stage changes that are occurring during this process.

Gene changes unique to Day 1 include genes implicated in tip/stalk specification such as KDR, CXCR4, JunB [18–20] as well as other elements of blood vessel development such as tubulogenesis (MMP14) [21] and proper growth control (MAP3K3, PGF, FYN) (**Figure 4a** and data not shown).

**Figure 4.**
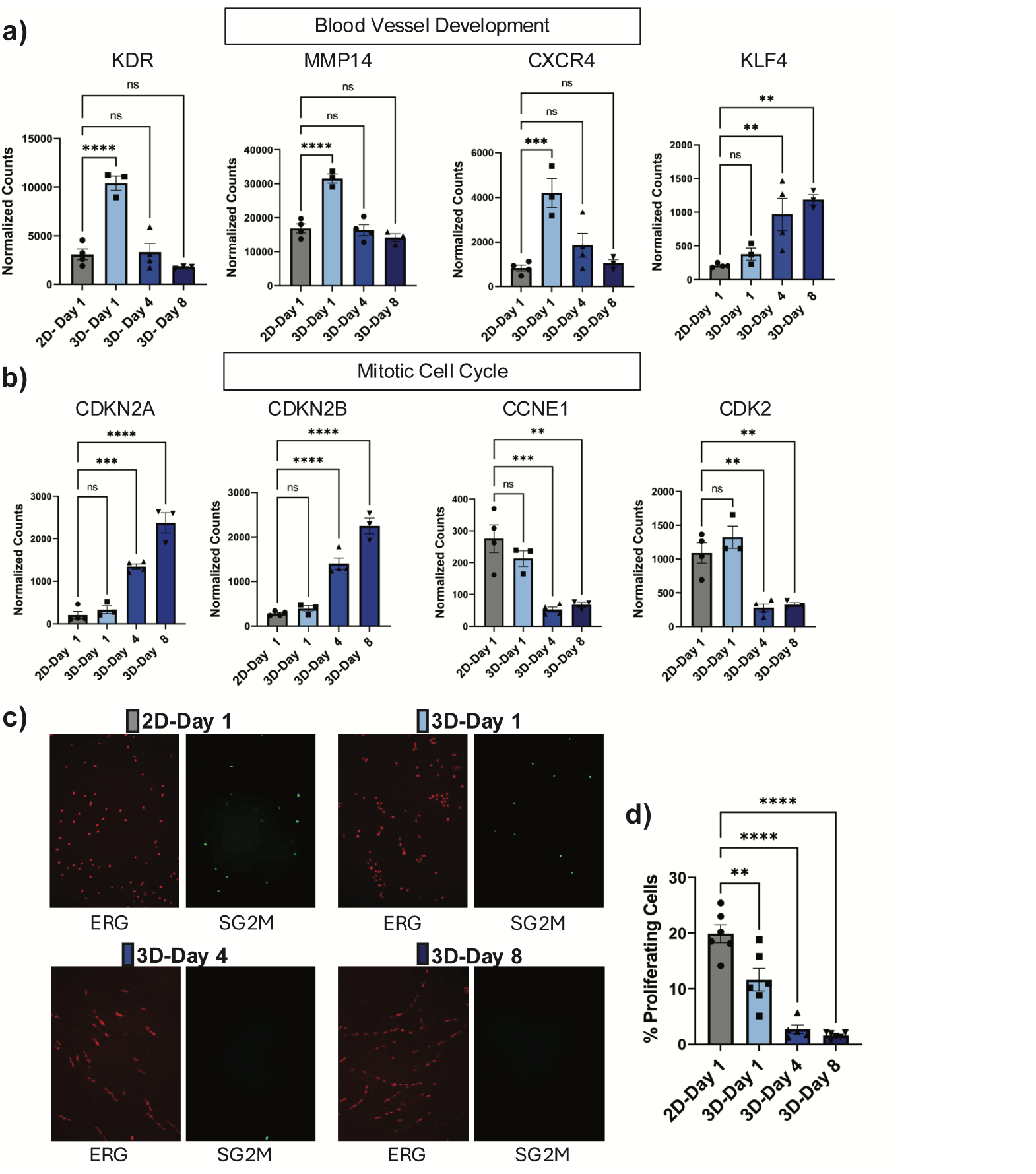
Genes involved in Blood Vessel Morphogenesis Show Distinct Expression Patterns Throughout Morphogenesis and Coordinate Cell Cycle Arrest. Quantification of selected genes in the blood vessel development GO term **(a)** and mitotic cell cycle **(b)**. Normalized raw counts are shown for the indicated time points. Significance is determined by One-way ANOVA, Dunnett’s (p<0.05). **c)** Representative images are shown of proliferating ECs expressing the S-G2-M proliferation reporter (green) and the endothelial specific nuclei stain ERG (red). **d)** Quantification of the percent of proliferating ECs (SG2M) per field of view, normalized to the number of EC nuclei present (ERG). Significance is determined by One-way ANOVA, Dunnett’s (p<0.05).

Gene Ontology analysis reveals the most robust enrichment of genes changes were in the mitotic cell cycle in both day 4 and day 8 samples. These changes are unique to day 8 compared to day 1 and include induction of several well-known cell cycle inhibitors such as, CDKN2A, CDKN2B, and CDKN1A (**Figure 4b** and Supporting data; Main 4c). This is accompanied by a significant concomitant downregulation of growth promoting cyclins and CDKs such as CCNE1, CDK2, CCNA1, CCNA2, and CDC20 (**Figure 4b** and Supporting data; Main 4c). These data strongly suggest a strong suppression of mitogenesis accompanies the differentiation we observe in the co-culture assay.

To confirm growth suppression *in situ* we used a S-G2-M proliferation reporter which fluoresces when a cell is progressing through the cell cycle, but which is rapidly degraded as the cell returns to G0/G1. We validated this reporter by comparing the number of cells showing positivity with the S-G2-M reporter compared to the well characterized thymidine analog, Edu.

Comparative assays were done in 2D HUVECs following serum starvation **(Supplemental Figure 4a and 4b)**, with nearly identical results at 24 hours post-stimulation. Using HUVECs infected with the proliferation reporter, 2D and 3D cultures were seeded and allowed to progress to various time points analogous to the RiboTag experiment. Consistent with the RNA-seq results, there was a significant decrease in the number of proliferating ECs as early as 1 day in the co-culture **(Figure 4c and 4d)**, with little to no remaining proliferation by day 8, despite the presence of a full complement of growth factors in the culture medium.

At day 8 we identified a number of genes from the *blood vessel development* GO term that were changing from day 1 to day 8 that are known to have links in NOTCH signaling. Due to the fundamental role NOTCH plays in proper angiogenesis control, we analyzed gene expression changes across the entire *NOTCH signaling pathway* term (GO: 0007219) **(Figure 5a).** We highlighted a subset of NOTCH genes that are known to have important roles in blood vessel development, NOTCH1, DLL4, JAG2, and SNAI2, that are dynamically regulated throughout the assay **(Figure 5b)**. In addition, genes representing several other ligand/receptor axes were uniquely changing at day 8 including TIE, TGFb, and PDGFb [22–26] which have all been implicated in blood vessel development and stabilization (Supporting data; Main 4c).

**Figure 5.**
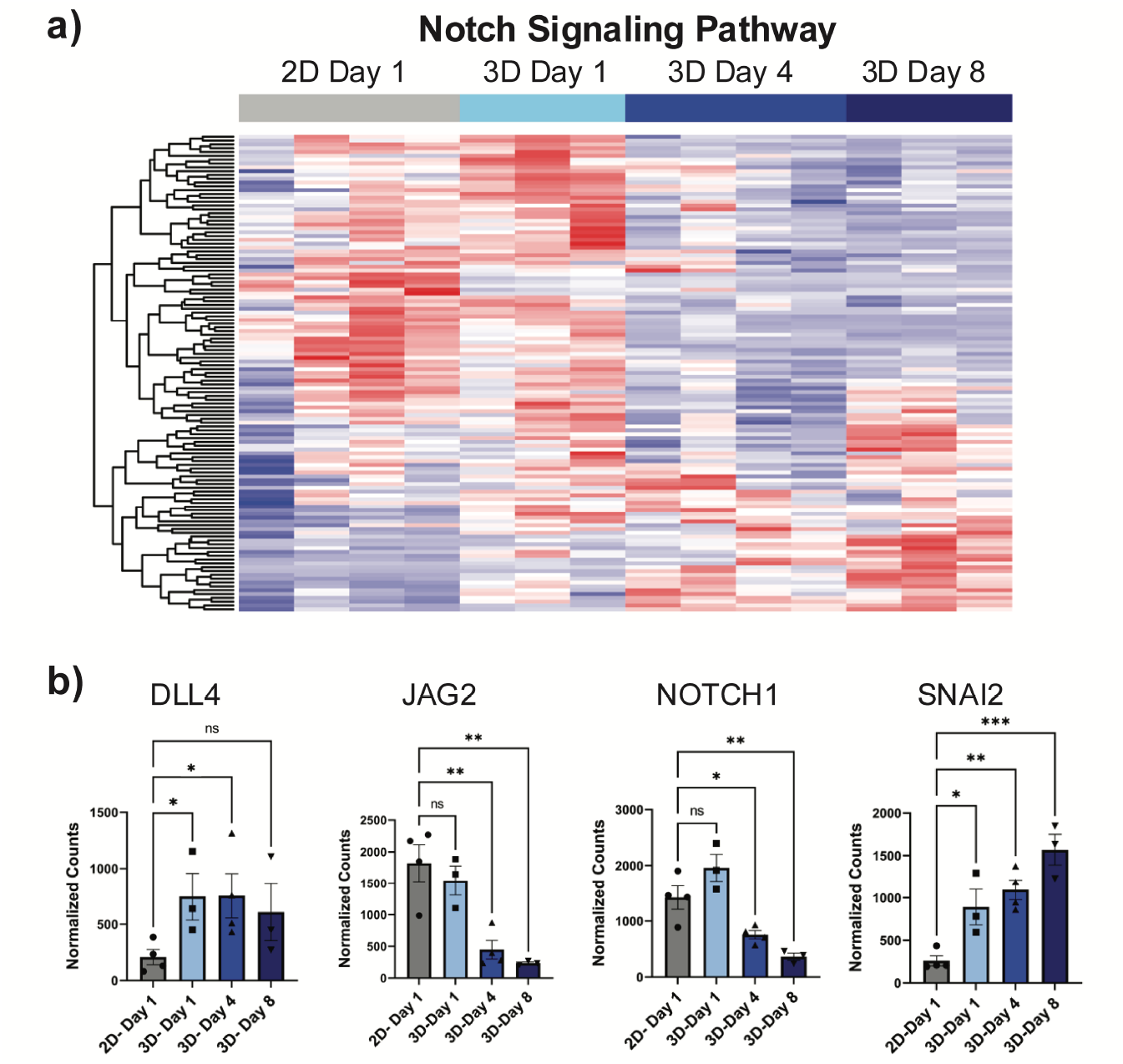
The NOTCH signaling pathway is dynamically regulated throughout morphogenesis. **a)** Heat map of genes from NOTCH signaling pathway GO term filtered for genes at baseline (2D and 3D day 1) are greater than 25 counts. **b)** Examples of genes from the NOTCH signaling pathway known to be important for tip/stalk regulation showing dynamic changes from early to late morphogenesis in the 3D culture. Significance determined for individual comparisons 2D vs. each 3D time point) by Student’s T-Test, * p Value <0.05. Individual comparisons were plotted on the same graph to show changes across time.

### Generation and validation of a morphogenesis gene signature

To identify genes that change in 3D culture, *independent of time point,* and to identify related changes in future transcriptional studies, we generated a gene enrichment signature of *in vitro* morphogenesis. This should provide a tool to readout gene expression changes important for EC differentiation, regardless of the time point in which the RNA is isolated. We generated this gene set by using multivariate analysis of 2D vs. all (3D day 1, day 4, and day 8) in Qlucore Software (release 3.1, https://qlucore.com/) and filtering for genes that have a Q-value cutoff less than 0.01. A heat map of the 365 genes generated by this analysis is shown in **Figure 6a**. We evaluated the biological processes enriched in this gene set and found that the top biological functions include angiogenesis, blood vessel morphogenesis and blood vessel development **(Figure 6b**), consistent with our goal to establish a time point independent signature of endothelial morphogenesis.

**Figure 6.**
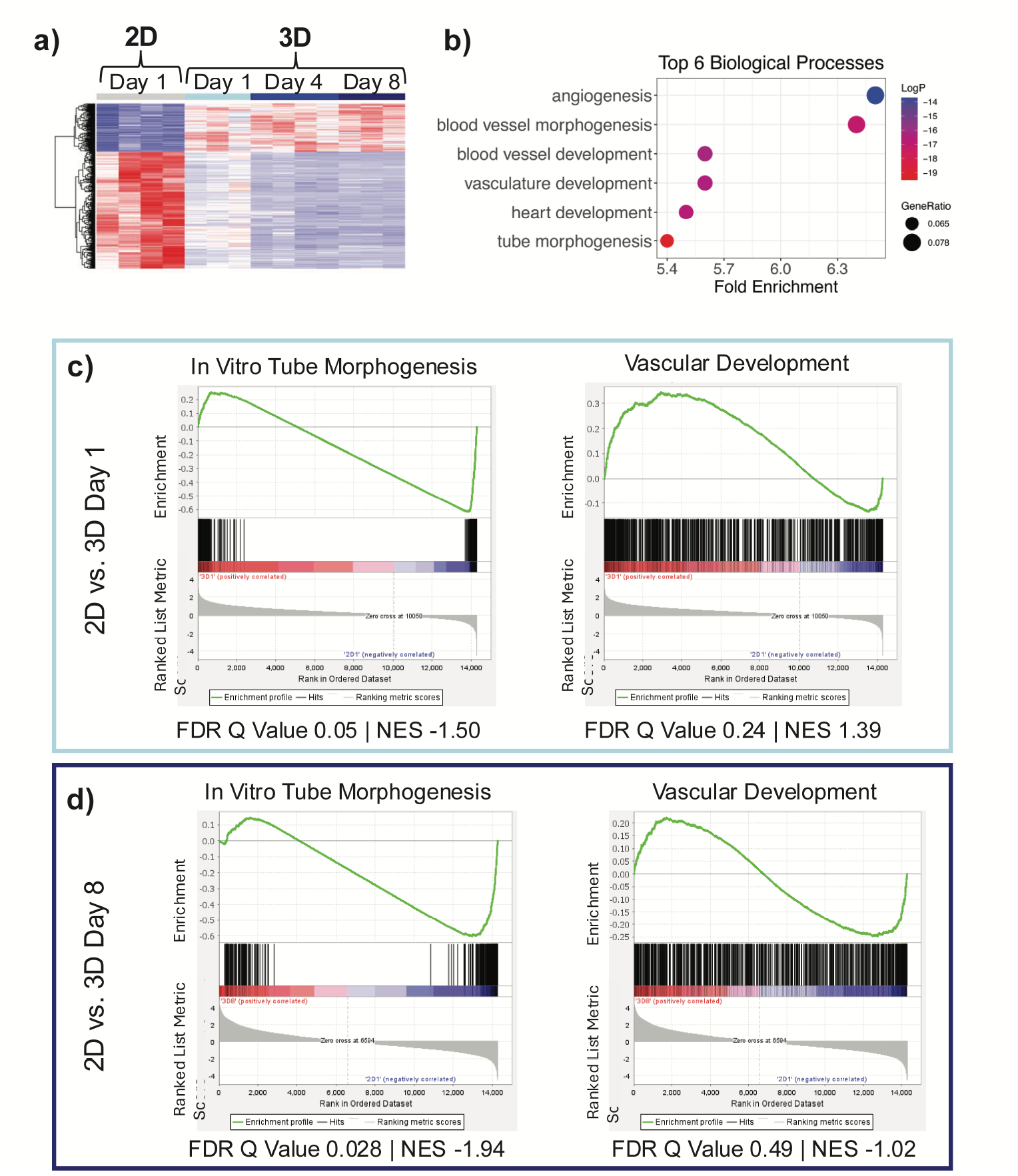
Generation of an in vitro morphogenesis gene signature. **a)** Heat map of genes in the morphogenesis gene signature that was generated by doing a multivariate analysis (2D vs. all (3D day 1, day 4, day 8) in Qlucore with a Q-value cut of 0.01. **b)** GO analysis from Metascape showing the top 6 biological processes enriched in the gene signature. **c)** Enrichment plot of GSEA analysis 2D and 3D day 1 using our *In Vitro Tube Morphogenesis* gene signature (left) or the existing Vascular Development signature (right). **d)** Enrichment plot of GSEA analysis of 2D and 3D day 8 using our *In Vitro Morphogenesis* gene signature (left)) or the existing *Vascular Development* signature (right).

To validate the quality of this signature, we compared the performance of this gene set (*In Vitro Tube Morphogenesis*) to the gene set enrichment analysis scores of an established vascular development gene set (*Vascular Development*, GO:0001944) which was the top performing match to our data by conventional GSEA analysis [27, 28] (using v2024 Hallmark gene set, https://www.gsea-msigdb.org/). Our generated signature showed superior enrichment and excellent false discovery metrics compared to the *Vascular Development* signature at both day 1 **(Figure 6c)** and Day 8 **(Figure 6d)**. These results suggest this gene set will be useful in detecting similar processes related to EC differentiation and tube morphogenesis in future transcriptomic studies, independent of whether it is early or late in the process.

## DISCUSSION

TRAP seq has been used to evaluate endothelial cell mRNA expression changes *in vivo* from a variety of organ systems, including the brain which is only 1-2% endothelial cells [13, 15, 17] thus we reasoned this would be useful as a technology to investigate changes in co-culture systems. Examining gene expression in complex 3D co-culture systems better mimics the *in vivo* environment and allows for the examination of perturbed states of morphogenesis. In this study, we found that TRAP seq can be applied to 3D multicellular cultures, including both the planar co-culture [4] and fibrin bead [5] assays. While there is a plethora of data on changes in EC gene expression using *in vitro* 2D cultures, these conditions, even at early culture time points, do not share transcriptional profiles of endothelial cells freshly isolated from umbilical cords [29]. In addition, it is well appreciated that there is a dynamic regulation of the endothelial cell transcriptional program by their surrounding microenvironment, as well as by physical forces such as flow [17, 29]. The ability to use TRAP seq in 3D co-cultures, and presumably even more complex organoid cultures, provides a methodology to isolate endothelial cells (or other cell types) from these complex differentiated environments rapidly and with limited artificial perturbation of the cells.

In our studies, gene expression changes were evident between 2D and 3D co-culture endothelial cells as early as 24 hours, with a strong enrichment of genes associated with vascular motility and development. Among the enriched set of genes, we saw increased expression of the well-known angiogenic chemokine receptor and EC tip cell gene, CXCR4. Neutralizing antibodies used against CXCR4 in angiogenesis assays in vitro, disrupt normal EC morphogenesis [30]. The ligand for this receptor, SDF1 (CXCL12), is also known to be critical for sprouting angiogenesis *in vitro* [31]. *In vivo*, CXCR4 has been shown to be specifically enriched in EC tip cells and when CXCR4 is inhibited in the developing retina, morphogenic defects occur [32]. We also found the receptor for VEGFA, KDR (also known as VEGFR2), to be upregulated at day1 when HUVECs are put into 3D co-culture. KDR has long been established to have a critical role in angiogenesis by regulating cell proliferation, differentiation, migration, and survival [33, 34]. Other genes known to be critical for angiogenesis also showed marked early changes in expression in angiogenesis including, UNC5B, DLL4, and MMP14, genes known to have roles in tip cell establishment and tube formation [18-20, 35, 36]. Many of these genes however reduced their expression over the course of eight days, while others more associated with vascular maturation and stability were increased, such as, KLF4 and various matrix proteins, underscoring the dynamic process that is observed in these cultures visually.

One of the major changes occurring from early to late morphogenesis was a vast downregulation of genes involved in cell cycle progression and proliferation. For the *mitotic cell cycle* gene ontology term alone, there was around 150 genes downregulated at both the day 4 and day 8 time points, which were not observed at day 1. Genes changing included well-known regulators of the G1-S transition such as the cell cycle inhibitors, CDKN2A and CDKN2B, which were increased, with a consistent decrease in the correspondingly regulated cyclins and CDKs such as, CCNE1, CDK2, CCNA1, and CCNA2. This powerful downregulation of proliferation was validated using a real-time S-G2-M proliferation reporter, which showed a dynamic and near complete reduction by day 4 in the co-culture. It is important to stress that this reduction is not due to the absence of mitogenic stimulation at the level of ligand availability, as these cultures contain a complete complement of growth factors and serum in the media. Rather it seems there is an active repression of the proliferative response that accompanies endothelial cell differentiation. This growth suppression likely coincides with establishment of junctional complexes, vessel stabilization, and maturation of the nascent tubules. ECs enter into a non-proliferative quiescent state in response to blood flow, fluid shear stress, pericyte recruitment, and formation of adherens junctions, all of which play a role in stabilizing the newly formed vessel *in vivo* [37]. To this end at the later time points, we observed an increase in KLF4, which has been shown to promote vascular integrity [38]. Moreover, we see changes in genes coding for junctional proteins, such as CLD5, which is known to be essential for barrier formation [39] as well as increased TEK (TIE2) and decreased in ANGPT2, suggesting increased capacity for TIE2 activation that is well described to enhance stabilization of the vasculature and modulate angiogenic signaling [22, 23, 40].

Collectively, our data presented here demonstrates that TRAP seq technology can be utilized in conjunction with complex co-culture models to study gene expression relevant to endothelial cell morphogenesis in both normal and presumably can be extended to investigate pathological contexts, such as those involved in vascular malformations. While these models are far from perfect, lacking flow, and proper investiture with stromal cells, they do mimic many of the hallmarks of early angiogenesis including sprouting, alignment, proliferative quiescence, and tubulogenesis. While mechanistic studies have been hampered in these systems by the cellular complexity, TRAP seq provides an attractive and approachable way to rapidly retrieve translating RNAs for expression analysis without the need for cellular purification and associated signaling perturbations. This should be a useful approach for refining molecular hypotheses, including the determination of how perturbations of genes critical for normal morphogenesis, such as those in the NOTCH signaling, might contribute to pathological angiogenesis. While our data analyzed the translatome of differentiating endothelial cells, this same technology can be used in other organoid systems such as those investigating epithelial cell differentiation [41], iPSC organoid formation [42], xenograft metastasis [43] and tumor microenvironment [44].

## Methods

### Cell culture

Human umbilical vein endothelial cells (HUVECS) from pooled donors were purchased from Lonza and PromoCell and cultured as we have previously described [6]. HUVECs are incubated in endothelial complete growth medium (ECGM2) purchased from PromoCell for the indicated times per each experiment. Primary human dermal fibroblasts (NHDFs) were purchased from PromoCell and cultured in 10% FBS DMEM.

### Plasmid construction

To construct the pHAGE-Rpl22-3XHA construct, the insert fragment (constructed by gene synthesis (Genewiz) and the pHAGE-IRES-tdTomato destination vector were digested with the NotI-HF (NEBioLabs #R3189S) and XhoI (NEBioLabs #R0146S) restriction enzymes. Following digestion, the Rpl22 fragment was ligated into the expression vector with the Instant Sticky-end Ligase Master Mix (NEBioLabs #M0370S). The final construct was then transformed into NEB Stable bacteria (NEBioLabs #C3040) to allow for plasmid DNA preparation.

### Viral infection and generation of stable cell lines

Lentiviruses were made by co transfection of the pHAGE-RPL22-3X-HA with the packaging plasmids pCMV-VCV-G and pCMV-delta) the Lipofectamine 3000 Transfection kit (Invitrogen L3000-001) into FS293 cells. After 24 hours media was replaced with 20% FBS (heat inactivated) MCDB131. High titer virus was harvested at 24 and 48 hours after 20% FBS MCDB131 addition. Low passage and sub-confluent HUVECs were infected with lentivirus for 5-6 hours with 4ug/mL polybrene. After infection, viral containing media is removed and replaced with ECGM2 and incubated for at least 24 hours. Infection efficiency was monitored using the expression of the td-Tomato (pHAGE-RPL22-3XHA) in stably infected cells. For 3D co-culture proliferation assays, cells are infected as described above, with the Fucci cell cycle vector, pRetroX-SG2M-Cyan (Takara bio, #631462).

### 3D Angiogenesis Assays

#### Planar Co-culture

This 3D *in vitro* angiogenesis assay was performed as previously described [4] with the following modification. Fibroblasts were seeded onto a gelatin coated 60mm dish 18 hours prior to endothelial cell seeding at a ratio 1:10 (HUVECS:Fibroblasts) and allowed to incubate for 1-8 days in ECGM2 (media changed every 2-3 days). Immediately prior to lysis for TRAP Seq, HUVECs were visualized and imaged with the internal td-Tomato fluorescent protein.

#### Fibrin bead assay

This was performed as previously described [5] and high-resolution images phase images were captured on Zeiss Axio Observer Z1 microscope.

### RNA Isolation, clean-up, and quantitation for TRAP Seq

Planar co-cultures are treated with an ice cold 1X PBS containing 100ug/mL cycloheximide wash and flash frozen. RNA lysates are then made with 400uL of polysome buffer for each 60mm dish of cells. Lysates are homogenized, and subsequently incubated with detergent (1% NP-40) for 20 minutes at 4 degrees. After incubation with the detergent, tubes are centrifuged for 10mins at 12,000 rpm at 4 degrees, and RNA containing supernatant is harvested. RNA was immunoprecipitated from planar co-cultures RNA lysate with Pierce anti-HA magnetic beads (Ref #88837) essentially as previously described [13]. RNA from the fibrin bead was isolated using a similar approach with slight modifications. Fibrin bead culture sizes were expanded to 100mm dishes. Cultures were flash frozen in liquid nitrogen prior to homogenization in 1mL of lysis buffer. Homogenized lysates from both assays are incubated with the anti-HA beads for 5-18 hours at 4 degrees. RNA was isolated from the magnetic beads with the Qiagen RNeasy Plus Micro Kit (Ref # 74034), the concentration determined using the Qubit RNA High Sensitivity Kit) Ref# Q32853 C), and the RIN score calculated using the Agilent Technologies RNA Pico Chips for use with the Agilent 2100 Bioanalyzer system (#5057-1513). Only Rin scores greater than 7 were used for further analysis.

### RNA-seq Data Processing and Differential Expression Analysis

RNA sequencing (RNA-seq) libraries were prepared by Azenta Life Sciences and sequenced on an Illumina NextSeq 500 platform, generating paired-end reads. Raw FASTQ files were trimmed using Trim Galore to remove adaptor sequences and low-quality bases. Alignment to the *Homo sapiens* reference genome (GRCh38/hg38) was performed using the *align* function from the Rsubread package [45] generating sorted BAM files. The mapping efficiency was assessed using the *propmapped* function [45]. Gene-level quantification was conducted using the *featureCounts* function in Rsubread, with annotation from the inbuilt hg38 gene model [45]. Only uniquely mapped, properly paired reads were counted.

Downstream analysis was performed in R using the edgeR and limma packages [46, 47]. Raw counts were converted into a DGEList object and annotated using the org.Hs.eg.db database from Bioconductor. Lowly expressed genes were filtered out using a threshold of ≥0.5 counts-per-million (CPM) in at least six libraries. Normalization for library size and composition bias was performed using the trimmed mean of M-values (TMM) method.

Differential expression analysis was carried out using the limma-voom pipeline. The *voom* function was used to model the mean-variance relationship and generate precision weights, which were then used in linear modeling via the *lmFit* function. Contrast matrices were defined for each comparison, and empirical Bayes moderation of variance estimates was applied using the *eBayes* function. Adjusted p-values were calculated using the Benjamini–Hochberg procedure to control the false discovery rate. Genes with an adjusted p-value < 0.05 were considered significantly differentially expressed.

### GSEA/Gene Signature

To generate the *in vitro* morphogenesis gene signature, we first performed a multivariate analysis of 2D culture vs. each 3D time point with a Q value cutoff of 0.01. This list of differentially expressed genes was used to create the “*in vitro* morphogenesis” hallmark gene set that is representative across various stages of morphogenesis. Enrichment of this set was validated in our own data and was compared to another hallmark gene set within MSigDB, “Vascular Development.” GSEA plots were generated using GSEA 4.3.3 a joint project of the Broad Institute and University of California San Diego (https://www.gsea-msigdb.org/gsea/index.jsp) [27, 48].

### Gene Ontology and R plots

Gene ontology analyses were completed using both the ShinyGO 0.82 hosted at South Dakota State University (https://bioinformatics.sdstate.edu/go/) [49] or with express analysis on Metascape (metascape.org) [50].For data generated by ShinyGO, the corresponding bubble plots shown in figures were generated by the ShinyGO. For those generated on Metascape, plots were regenerated to match the format of those by ShinyGO using the R package ggplot2 (R version 2024.04.2+764). Heatmaps were also generated in R using the ggplot2 package [51].

### Statistical analysis

All statistical analysis and graphs were made in GraphPad Prism or in R. Experiments were conducted using at least 3 (4 for most) independent pools of primary HUVECs from two different commercial vendors, each pool representing at least three donors of mixed races and sexes.

Each ‘n’ represents an independent sample. Statistical significance was performed by two-tailed one- or two-sample unpaired Student’s t-test, or one-way ANOVA with Dunnett’s post-hoc. All sequencing P-values are adjusted for multiple comparisons (Benjamini–Hochberg method, FDR). Details of the statistical analysis are present in the figure legends.

## Supporting information

Full page main figures

## Data Availability

All analysis for the assessment of differential expression (TRAP seq data) was processed in R. R Code files are available upon request. RNA seq datasets are available from GEO using the following identifier: Planar co-culture morphogenesis (GSE296693). Upon publication the GSEA signature will be submitted to mSigDB for consideration. The plasmid pHAGE-RPL22-3X-HA will be deposited in ADDGENE. Any additional data or methods is available from the authors upon request to the corresponding author at.

