## Supplementary material for "TRAP seq in 3D Angiogenesis Assays Reveals a Distinct Endothelial Translatome Associated with Early and Late Stages of Morphogenesis": Full page main figures

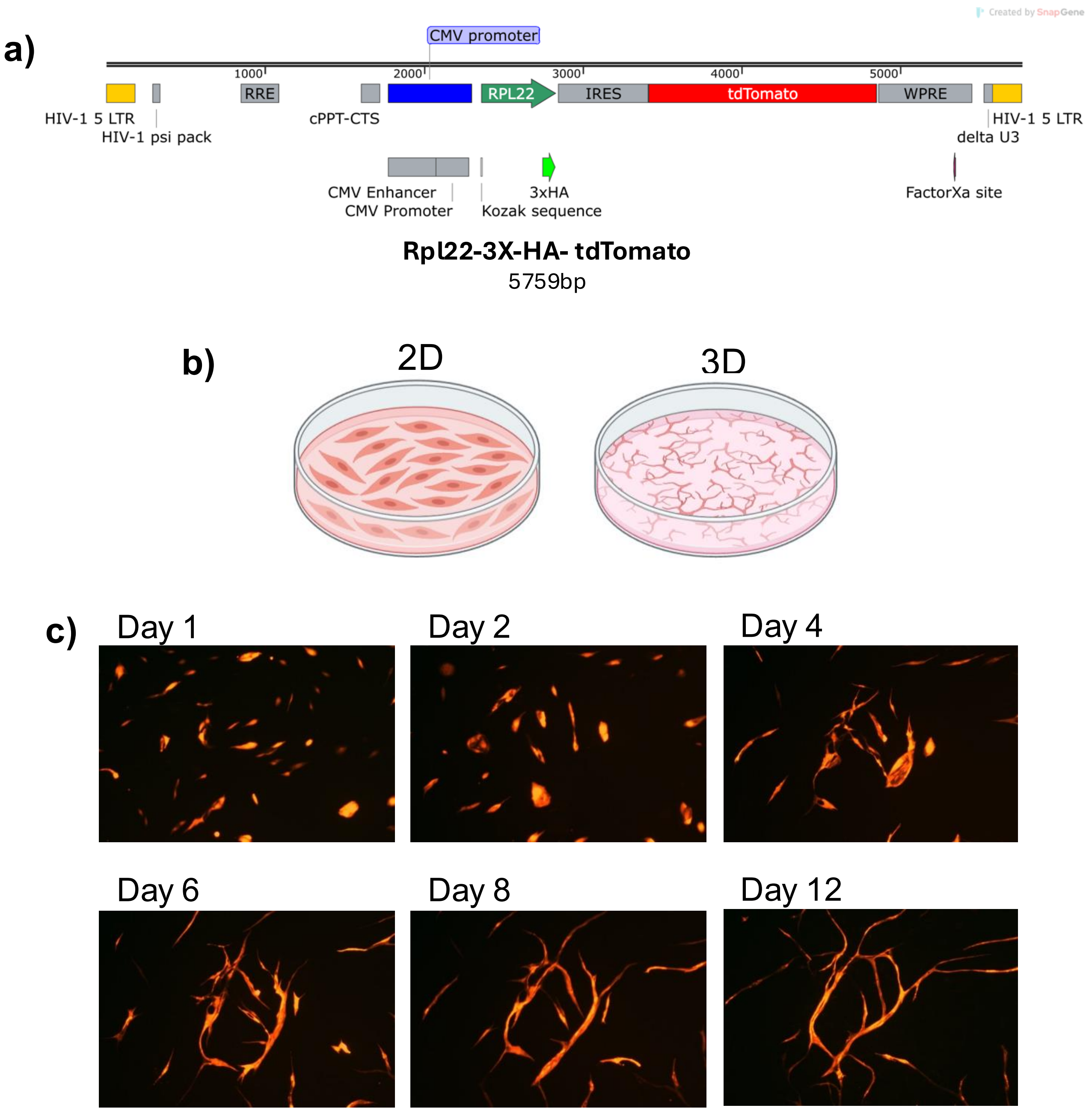


**Figure 1: Generation of a lentiviral vector harboring 3X-HA-Rpl22 and its use in a 3D co-culture of HUVECs.**
**a)** Map of 3XHA-Rpl22-tdTomato insert used in pHAGE lentiviral backbone for use in HUVECs to be subjected to TRAP seq. **b)** Diagram depicting morphological differences in 2D culture of HUVECs compared to 3D culture of HUVECs cultured on top of a confluent monolayer of fibroblasts in a planar co-culture morphogenesis assay **c)** HUVECs infected with 3X-HA-Rpl22-tdTomato are co-cultured with primary fibroblasts for 12 days in the planar co-culture assay. Images are taken at 5X at the indicated time points and then equally cropped via ImageJ to highlight anastomoses. Images are representative of 4 independent experiments.

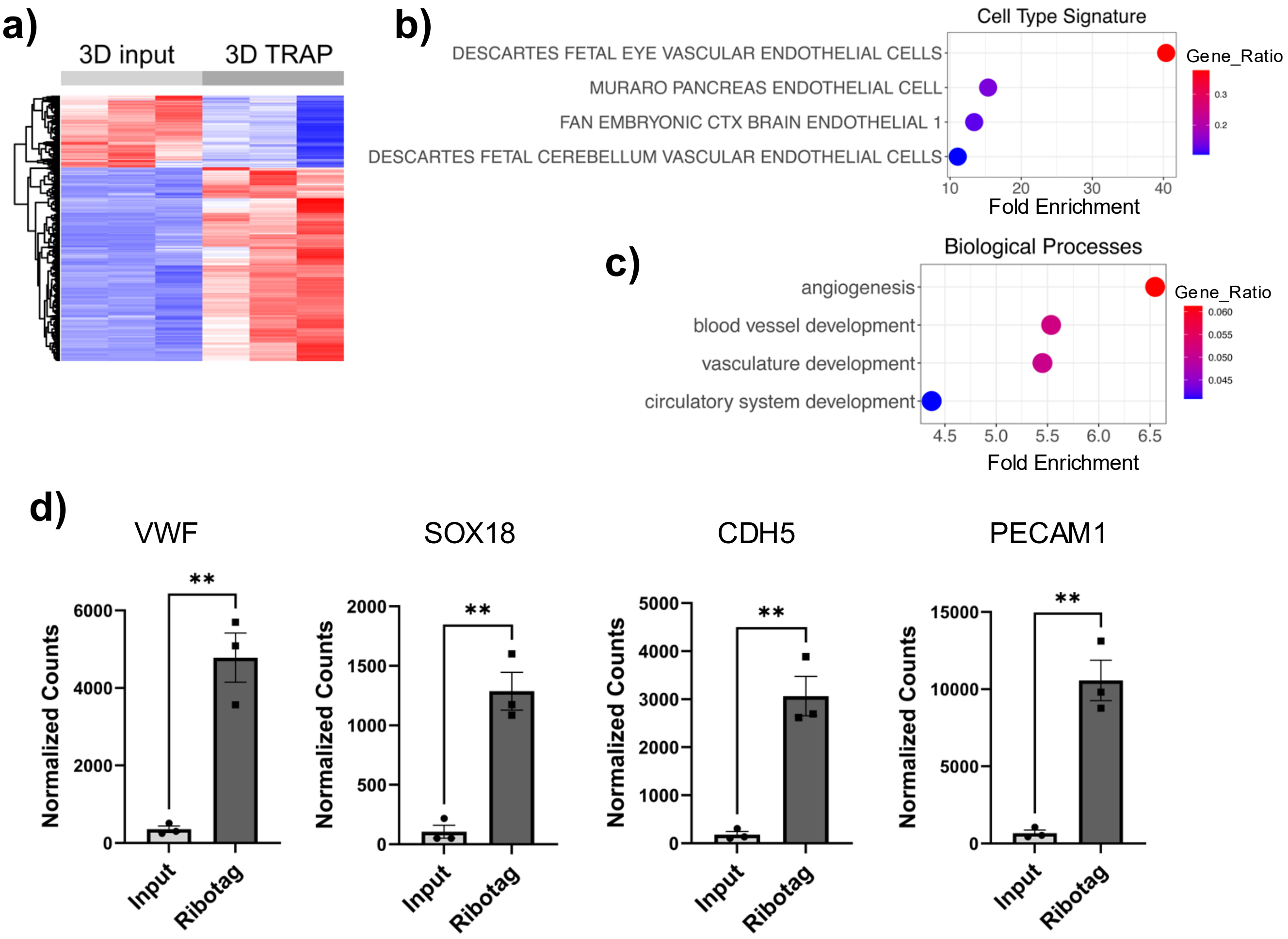


**Figure 2: RiboTag provides significant enrichment of endothelial cell transcripts from planar co-culture.**
**a)** Heat map of differentially expressed genes between input and TRAP-seq samples filtered for genes changing 1-fold with $p<0.05$. **b)** TRAP DEGs changing increasing more than 3-fold ($p <0.05$), were evaluated against cell-type signatures in MSigDB, with the top terms shown. **c)** Top GO terms in biological processes following analysis of upregulated DEGs changing 3-fold ($p<0.05$). **d)** Normalized raw counts showing enrichment of classic EC marker genes VWF, SOX18, CDH5, and PECAM1 in TRAP samples significance determined by student's t-test, $p<0.05$.

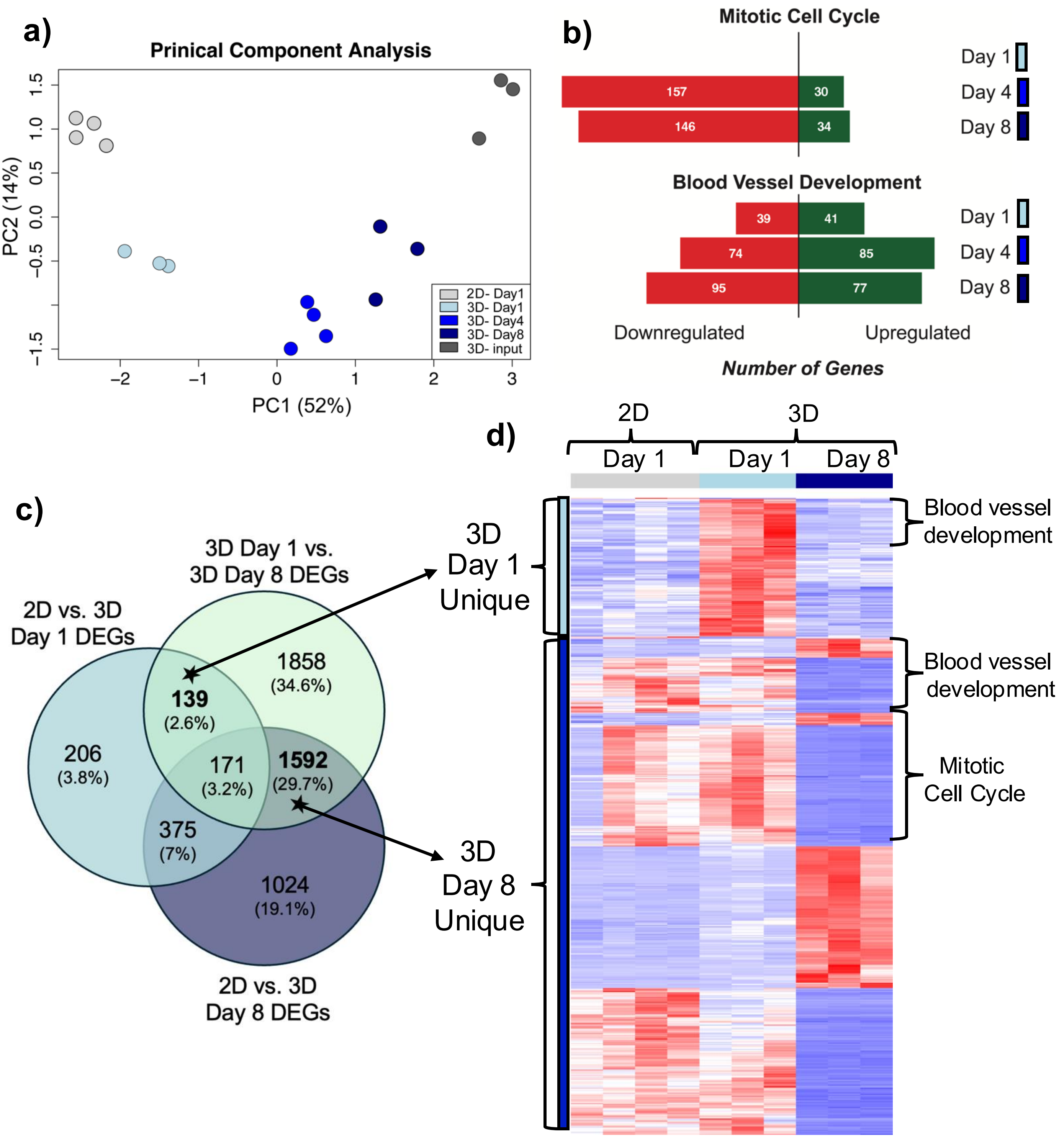


**Figure 3. RiboTag reveals distinct biological processes involved with normal vascular morphogenesis.**

**a)** PCA plot of groups included in TRAP seq analysis. **b)** Top GO terms from Metascape analysis (p-value <0.05 and logFC >1 and logFC< -1) of DEGs between 2D day 1 and 3D day 1, or 3D day 4, or 3D day 8. The number of individual gene hits for that GO term that are upregulated are shown in green and downregulated are shown in red. **c)** Venn diagram from set analysis between 2D vs. 3D day 1 (light blue), 2D vs. 3D day 8 (dark blue), and 3D day 1 vs. 3D day 8 (gray). This comparison allowed for the selection of unique genes that are important for 3D culture at early or late time points **d)** Genes unique to days 1 and 8 were combined and clustered to represent by a Heatmap. Genes derived from the blood vessel development and mitotic cell cycle GO term are highlighted. Day 1 unique genes are denoted by the light blue bars, and the day 8 unique genes by the dark blue bars (DEGs with Log FC +/- 1, p-value <0.05).

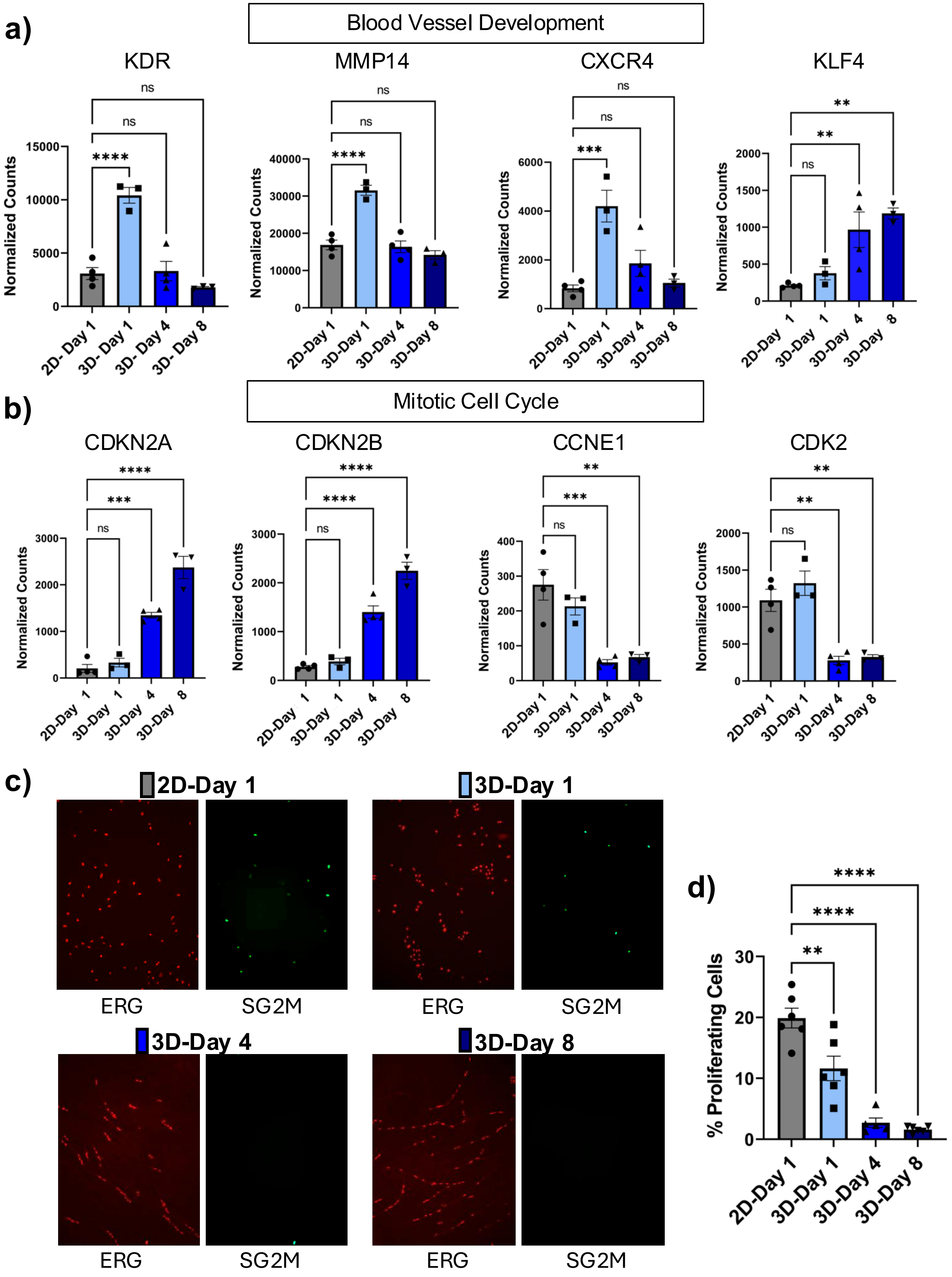

a)
Blood Vessel Development
KDR
MMP14
CXCR4
KLF4
Normalized Counts
ns
****
**
2D- Day 1
3D- Day 1
3D- Day 4
3D- Day 8
b)
Mitotic Cell Cycle
CDKN2A
CDKN2B
CCNE1
CDK2
***
2D-Day 1
3D-Day 1
3D-Day 4
3D-Day 8
c)
2D-Day 1
3D-Day 1
3D-Day 4
3D-Day 8
ERG
SG2M
d)
% Proliferating Cells

**Figure 4. Genes involved in Blood Vessel Morphogenesis Show Distinct Expression Patterns Throughout Morphogenesis and Coordinate Cell Cycle Arrest.**
Quantification of selected genes in the blood vessel development GO term **(a)** and mitotic cell cycle **(b)**. Normalized raw counts are shown for the indicated time points. Significance is determined by One-way ANOVA, Dunnett's ($p<0.05$). **c)** Representative images are shown of ECs expressing the S-G2-M proliferation reporter to enhance contrast. The nuclei of cells proliferating are shown in white. **d)** Quantification of the number of proliferating cells per field of view. Significance is determined by One-way ANOVA, Dunnett's ($p<0.05$).

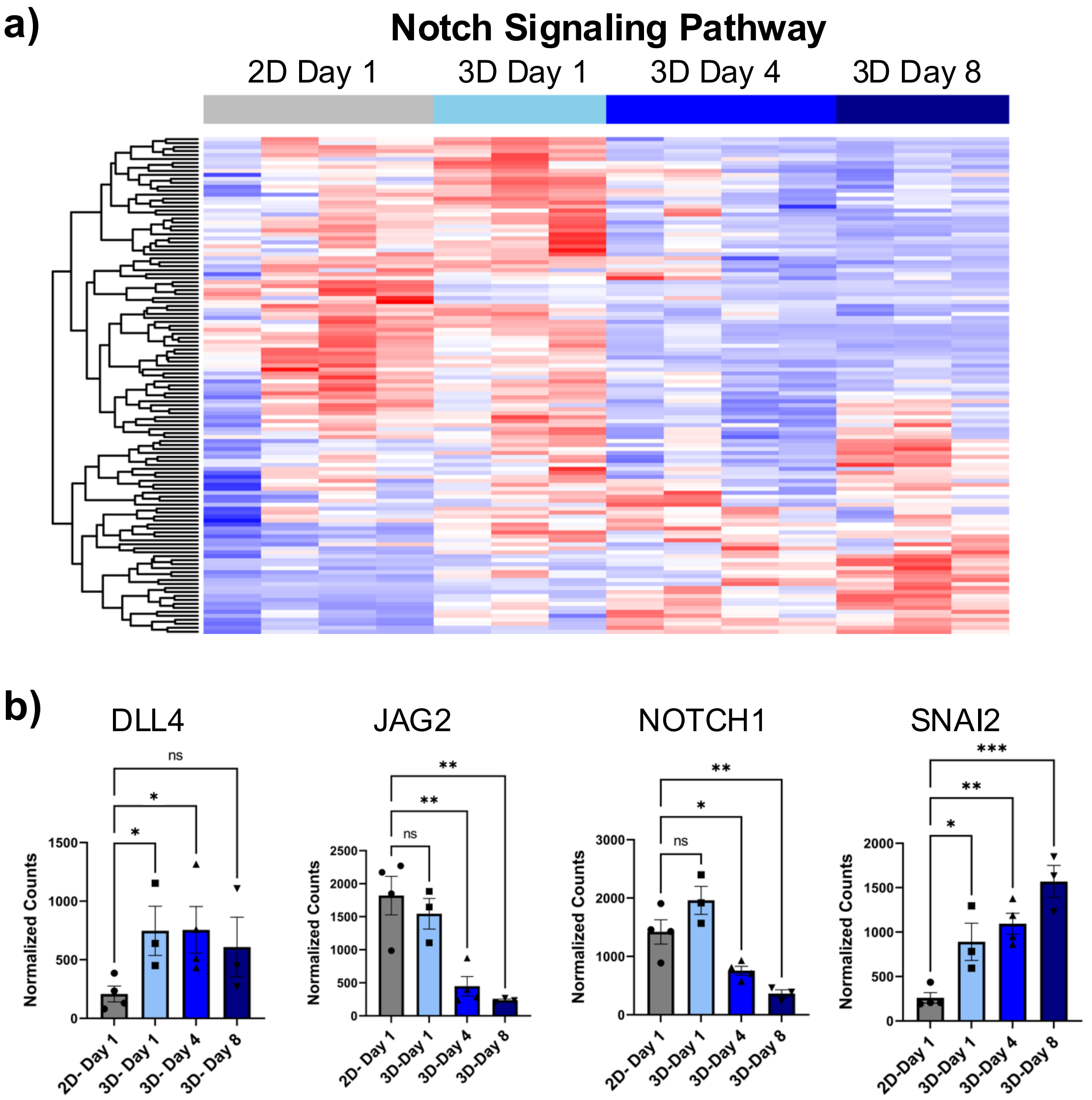


**Figure 5. The NOTCH signaling pathway is dynamically regulated throughout morphogenesis.**
**a)** Heat map of genes from NOTCH signaling pathway GO term filtered for genes at baseline (2D and 3D day 1) are greater than 25 counts. **b)** Examples of genes from the NOTCH signaling pathway known to be important for tip/stalk regulation showing dynamic changes from early to late morphogenesis in the 3D culture. Significance determined for individual comparisons 2D vs. each 3D time point) by Student's T-Test, * p Value <0.05. Individual comparisons were plotted on the same graph to show changes across time.

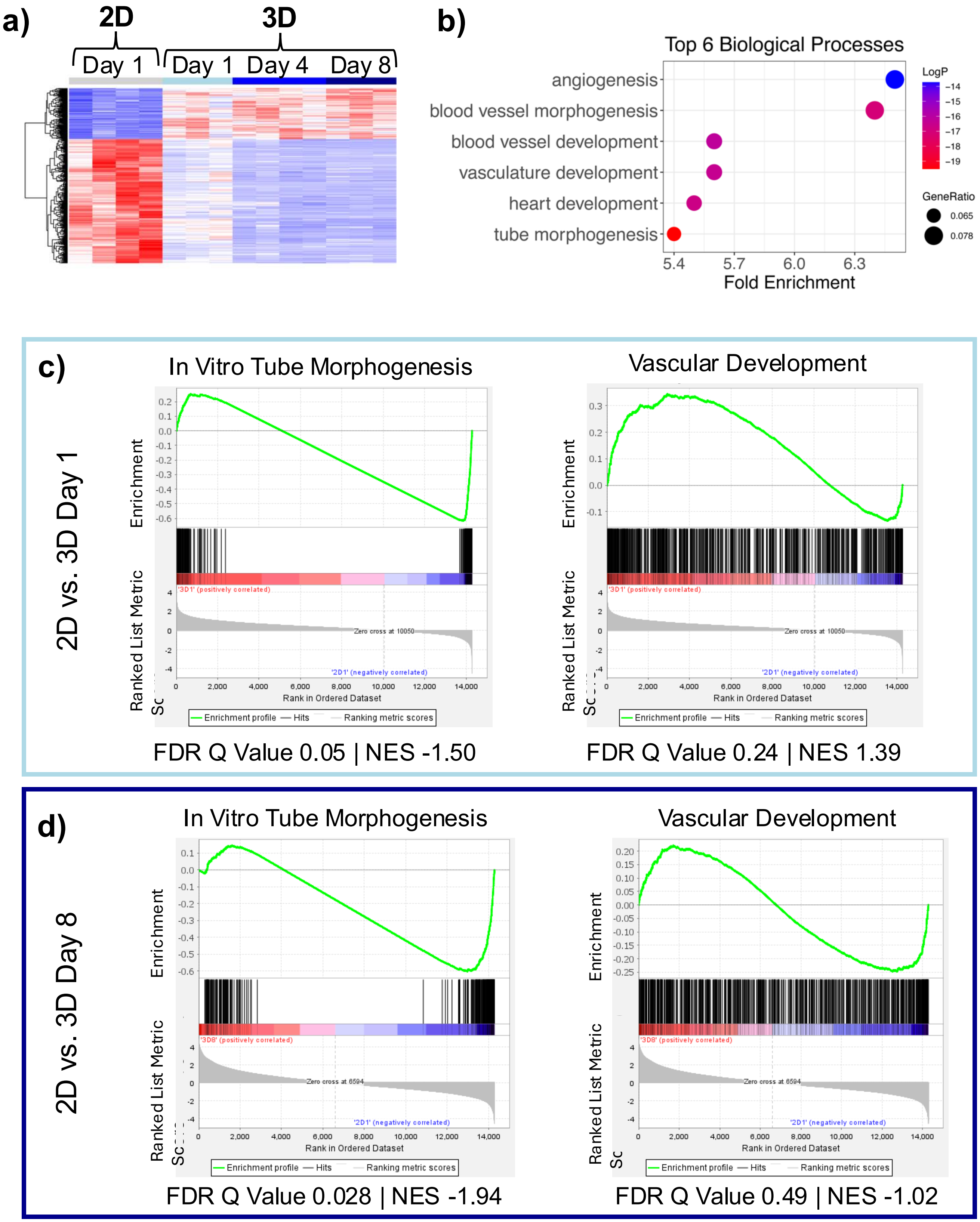
a)
2D
3D
Day 1
Day 1
Day 4
Day 8
b)
Top 6 Biological Processes
angiogenesis
blood vessel morphogenesis
blood vessel development
vasculature development
heart development
tube morphogenesis
5.4
5.7
6.0
6.3
Fold Enrichment
LogP
−14
−15
−16
−17
−18
−19
GeneRatio
0.065
0.078
c)
2D vs. 3D Day 1
In Vitro Tube Morphogenesis
Vascular Development
Enrichment
Ranked List Metric
Rank in Ordered Dataset
'3D1' (positively correlated)
'2D1' (negatively correlated)
Zero cross at 10050
Enrichment profile
Hits
Ranking metric scores
FDR Q Value 0.05 | NES -1.50
FDR Q Value 0.24 | NES 1.39
d)
2D vs. 3D Day 8
In Vitro Tube Morphogenesis
Vascular Development
'3D8' (positively correlated)
'2D1' (negatively correlated)
Zero cross at 6594
FDR Q Value 0.028 | NES -1.94
FDR Q Value 0.49 | NES -1.02

**Figure 6. Generation of an in vitro morphogenesis gene signature.**
**a)** Heat map of genes in the morphogenesis gene signature that was generated by doing a multivariate analysis (2D vs. all (3D day 1, day 4, day 8) in Qlucore with a Q-value cut of 0.01. **b)** GO analysis from Metascape showing the top 6 biological processes enriched in the gene signature. **c)** Enrichment plot of GSEA analysis 2D and 3D day 1 using our *In Vitro Morphogenesis* gene signature (left) or the existing Vascular Development signature (right). **d)** Enrichment plot of GSEA analysis of 2D and 3D day 8 using our *In Vitro Morphogenesis* gene signature (left)) or the existing *Vascular Development* signature (right).
