## Supplemental Figures 1-4 for "TRAP seq in 3D Angiogenesis Assays Reveals a Distinct Endothelial Translatome Associated with Early and Late Stages of Morphogenesis"

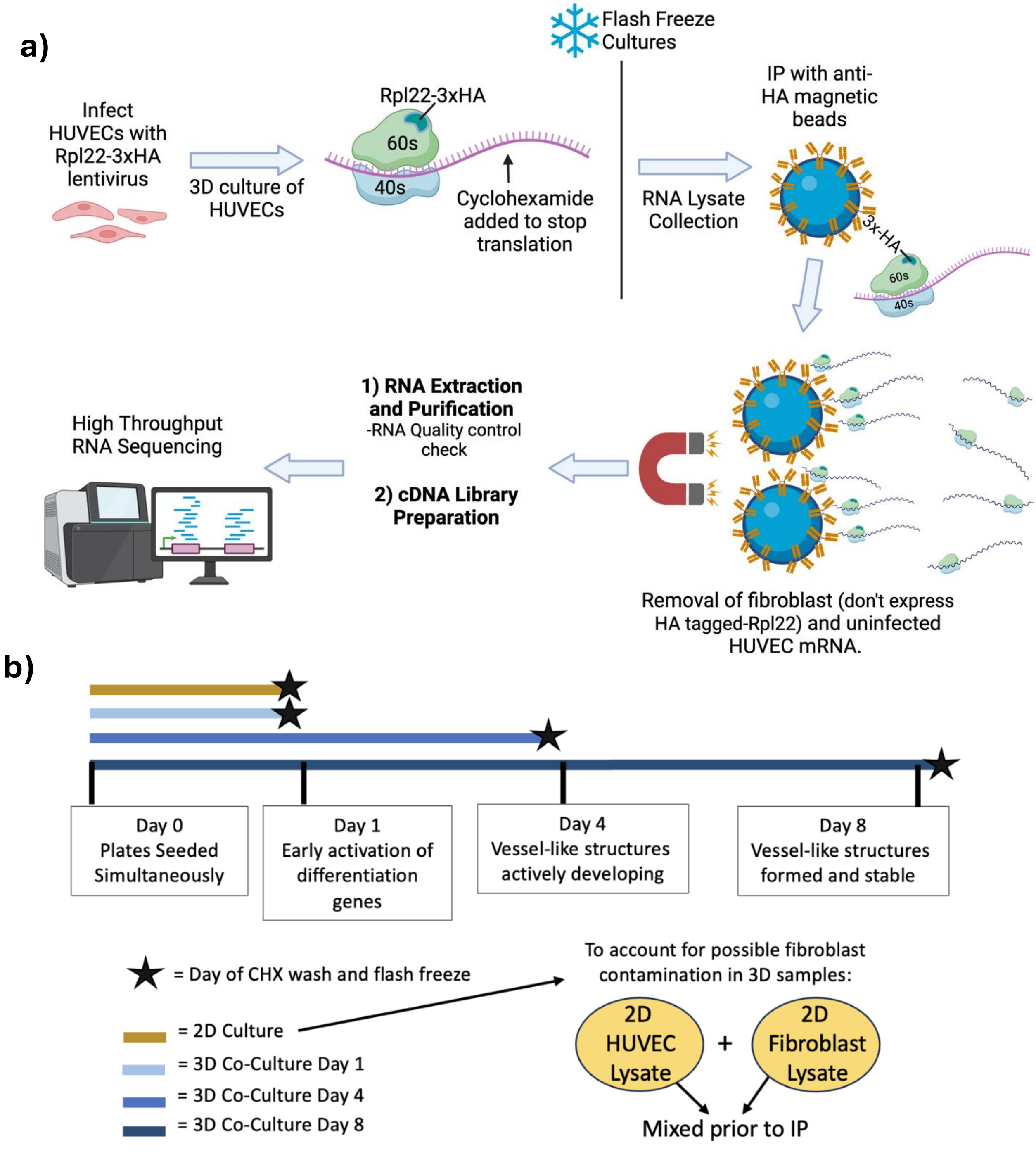


**Supplement figure 1: TRAP diagram and Timeline for Sample Collection for Planar Co-Culture TRAP Seq analysis**

**a)** Diagram showing the process of TRAP-Seq. HUVECs are infected with lentivirus coding for HA-Rpl22 and seeded into 3D co-culture. When the cultures are ready for termination, the cells are treated with a PBS/cycloheximide wash to halt translation. Cultures are then flash frozen to preserve HUVECs in their current differentiated state as they are undergoing morphogenesis. Lysis is initiated on ice and then immunoprecipitated with anti-HA magnetic beads. mRNA bound to the magnetic beads is isolated and analyzed for quality control parameters prior to being sent for high throughput RNA sequencing. **b)** Diagram showing the timeline for seeding of each lot of HUVECs for the co-culture and the subsequent termination of the culture (PBS+CHX wash and flash freeze). All cells from one lot are seeded simultaneously for all time points. To account for any fibroblast contamination that might be present in 3D cultures of HUVECs in the 2D culture condition, a common lysate is generated from separately cultured 2D endothelial and fibroblasts in numbers identical to those plated in the co-culture.

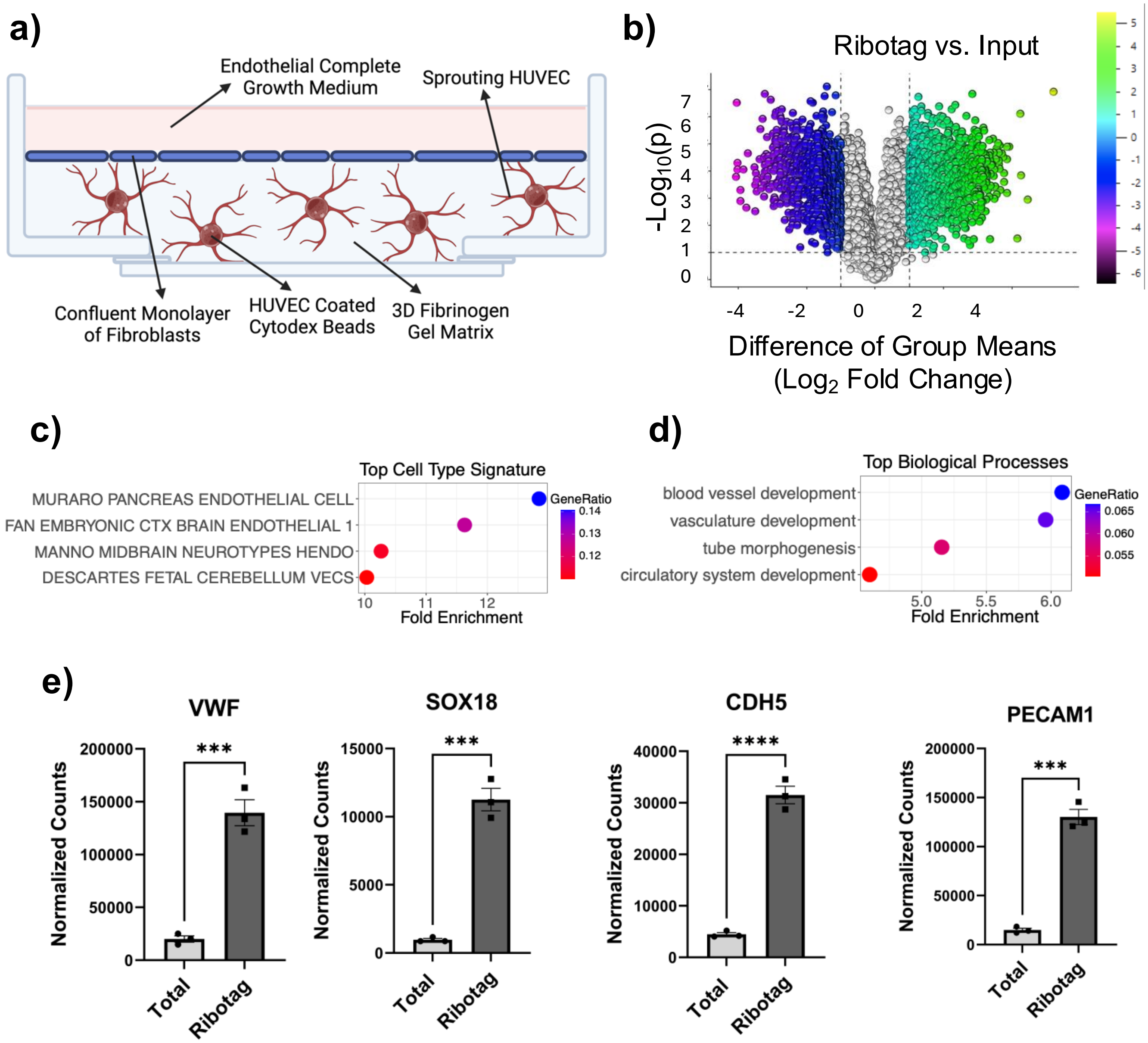


**Supplemental Figure 2. TRAP Validation in the 3D Bead Assay.**

**a)** Diagram displaying components of the 3D fibrin bead morphogenesis assay where HUVECs differentiate into a 3D fibrin matrix when co-cultured with fibroblasts. HUVECs are coated onto Cytodex beads and then embedded into a fibrinogen gel matrix. After the gel has clotted, fibroblasts are seeded in a 1:10 ratio on top of the gel, and ECGM is added. **b)** Volcano plot of DEGs between input vs. IP samples from the fibrin bead assay of HUVECs expressing Rpl22-3XHA. **c)** Top 4 cell types enriched in TRAP DEGs. **d)** Top 4 upregulated GO terms from. **e)** Enrichment of classic EC genes VWF, SOX18, CDH5, and PECAM1 in IP samples (normalized counts) significance determined by student’s t-test, $p<0.05$.

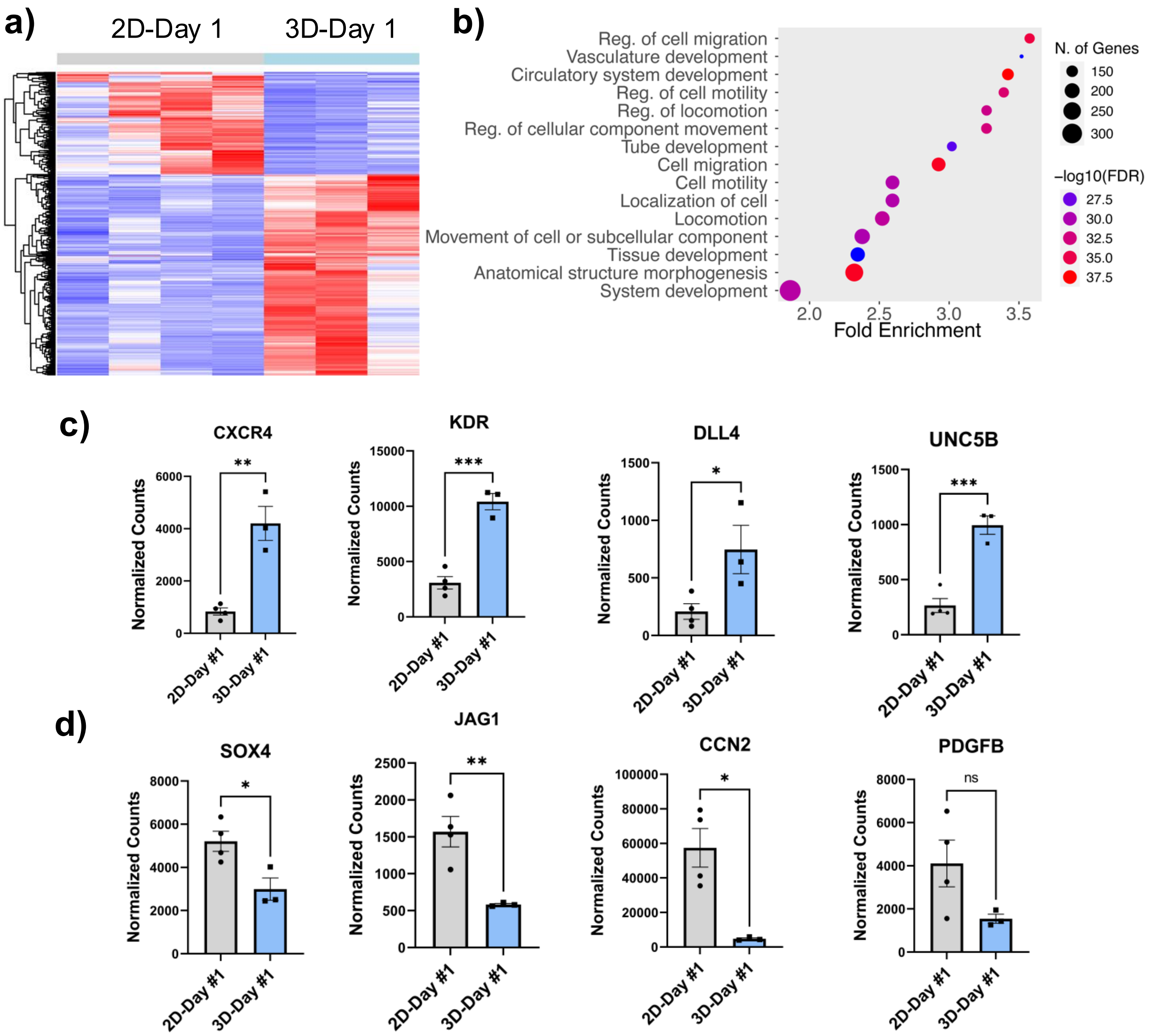


**Supplemental Figure 3: HUVECs cultured in the planar co-culture 3D organoid model results in changes to morphogenic gene expression compared to 2D culture.**

**a)** Heat map of top 578 DEGs with a p-value <0.01 and FC > +/- 2. **b)** Top 10 biological processes GO terms that change with 3D culture as determined by ShinyGO analysis (p-value <0.05 and logFC >2 and logFC< -2. **c)** Normalized counts of upregulated genes in TRAP samples, CXCR4, KDR, DLL4, and UNC5B. **d)** Normalized counts of downregulated genes in TRAP samples. Significance determined by student's t-test, p<0.05.

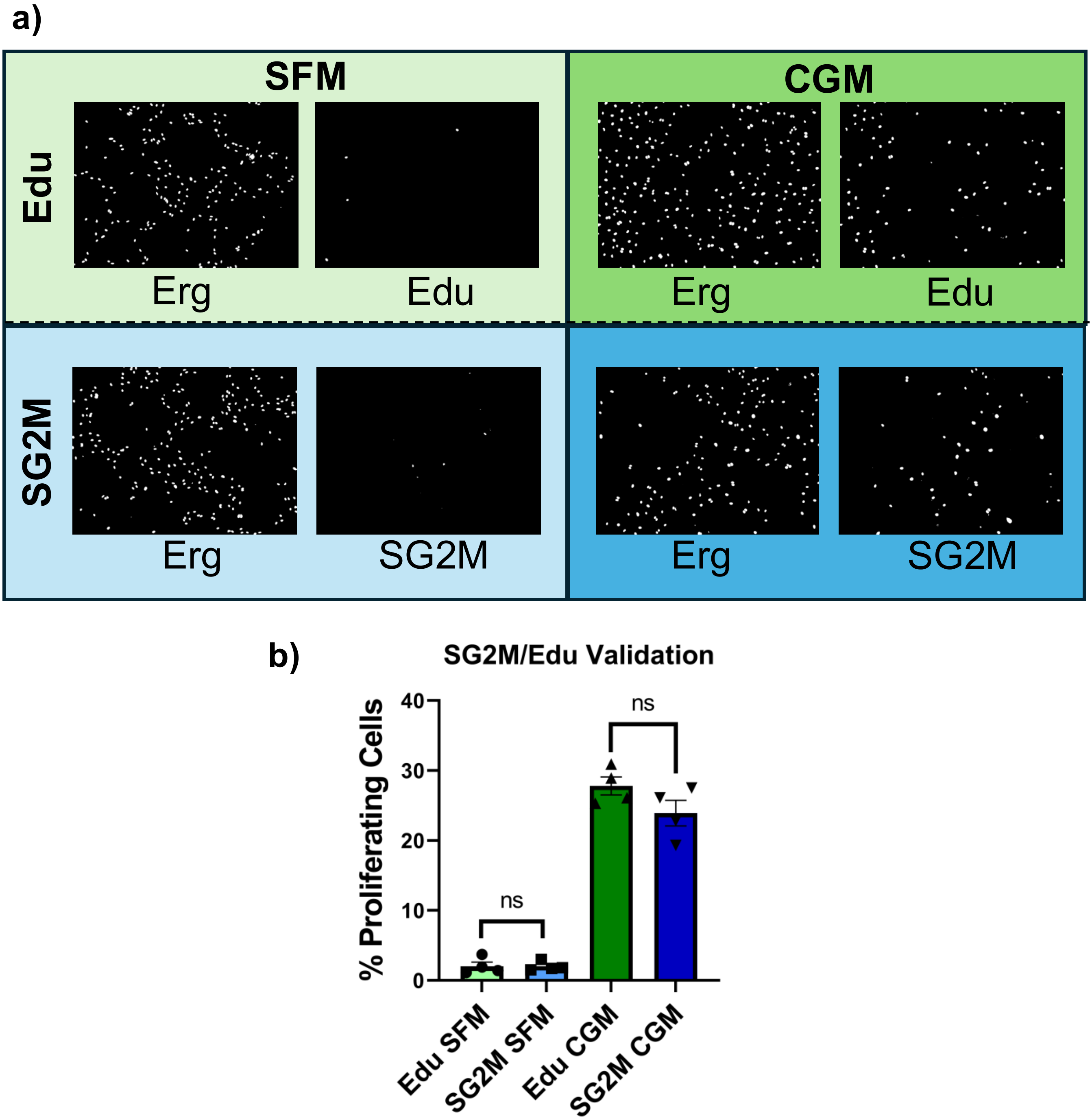


**Supplemental Figure 4: Validation of the SG2M Proliferation Reporter.**
**a)** Representative images of cells incubated with Edu (green/top) or infected with the SG2M proliferation reporter (blue/bottom). Cells were incubated in either complete growth medium (Left) or serum free medium (right) for 24 hours. HUVEC nuclei were stained with Erg to determine the total number of HUVECs. This value was used to determine the percent of proliferation cells for each condition. **b)** Quantification of images in **A** (N=4). Significance determined by one-way ANOVA, Dunnett's test ($p < 0.05$).
